# Lifelong olfactory deprivation-dependent cortical reorganization restricted to orbitofrontal cortex

**DOI:** 10.1101/2023.01.25.525521

**Authors:** Moa G. Peter, Fahimeh Darki, Evelina Thunell, Gustav Mårtensson, Elbrich M. Postma, Sanne Boesveldt, Eric Westman, Johan N. Lundström

## Abstract

Prolonged sensory deprivation has repeatedly been linked to cortical reorganization. We recently demonstrated that individuals with congenital anosmia (CA, complete olfactory deprivation since birth) have seemingly normal morphology in piriform (olfactory) cortex despite profound morphological deviations in the orbitofrontal cortex (OFC), a finding contradictory to both the known effects of blindness on visual cortex and to the sparse literature on brain morphology in anosmia. To establish whether these unexpected findings reflect the true brain morphology in CA, we first performed a direct replication of our previous study. Individuals with CA (n=30) were compared to age and sex matched controls (n=30) using voxel- and surface-based morphometry. The replication results were near identical to the original study: bilateral clusters of group differences in the OFC, including CA atrophy around the olfactory sulci and volume increases in the medial orbital gyri. Importantly, no group differences in piriform cortex were detected. Subsequently, to assess any subtle patterns of group differences not detectable by our mass-univariate analysis, we explored the data from a multivariate perspective. Combining the newly collected data with data from the replicated study (CA=49, Control=49) we performed support vector machine classification based on gray matter volume. In line with the mass-univariate analyses, the multivariate analysis could accurately differentiate between the groups in bilateral OFC (75-96% accuracy), whereas the classification accuracy in piriform cortex was at chance level. Our results suggest that despite lifelong olfactory deprivation, piriform (olfactory) cortex is morphologically unaltered and the morphological deviations in CA are confined to the OFC.

## 1. INTRODUCTION

Sensory deprivation is commonly linked to morphological and functional reorganization in, but not limited to, core processing regions of the deprived sensory modality (Lee and Whitt, 2015; Noppeney, 2007; Voss, 2019). However, in sharp contrast to prior publications (Frasnelli et al., 2013; Karstensen et al., 2018) and to our own a priori expectations, we recently demonstrated a complete lack of morphological abnormalities in piriform (olfactory) cortex in a comparably large sample of adult individuals with congenital anosmia (CA, complete olfactory sensory deprivation since birth; (Peter et al., 2020). This striking absence of altered morphology in piriform cortex, despite lifelong sensory loss, was accompanied by clear bilateral abnormalities in the orbitofrontal cortex (OFC), a region vital for higher-order olfactory processing (Gottfried and Zald, 2005; Lundström et al., 2011). Here, we aim to establish whether these unexpected results accurately reflect the brain morphology in individuals with CA by first replicating the analysis of our previous publication on a newly collected data set and subsequently applying a potentially more sensitive multivariate analysis approach on both the previously published and newly collected data, thereby enabling detection of potential morphological deviations not detectable by the previously used mass-univariate analysis methods.

The piriform cortex is unquestionably important for human olfactory processing: it is the main recipient of input from the olfactory bulbs (Gottfried, 2006; Lundström et al., 2011) and demonstrates the highest degree of odor activation in neuroimaging studies (Seubert et al., 2013). It is commonly referred to as primary olfactory cortex (along with the other regions receiving direct bulbar input (Gottfried, 2006; Zhou et al., 2019; however see Haberly, 2001) and has been linked to a multitude of functions, from representation of the chemical constitution of odors to odor category representation (Freiherr, 2017; Gottfried, 2010) with predictive and attentional modulation (Zelano et al., 2011, 2005). Olfactory sensory deprivation has been linked to altered morphology in piriform cortex as well as in other olfactory processing regions (for reviews, see Boesveldt et al., 2017; Manan et al., 2022; Reichert and Schöpf, 2018), analogous to the demonstrated morphological reorganization of early visual processing regions in blind individuals (Jiang et al., 2015, 2009; Noppeney et al., 2005; Park et al., 2009; Ptito et al., 2008). However, the literature on brain morphology in anosmia in general, and congenital anosmia in particular, is sparse and there is little overlap of results between studies. Specifically, there are to the best of our knowledge only three studies statistically assessing whole brain gray matter morphology differences between individuals with CA and healthy controls: Frasnelli et al. (2013), Karstensen et al. (2018), and from our lab, Peter et al. (2020). All three studies demonstrate morphological group differences in olfactory-related regions such as the OFC, but of particular interest is the fact that the first two studies indicate group differences in piriform cortex, whereas, contradictory to the stated hypothesis, we were unable to replicate these findings despite a larger sample (Peter et al., 2020). In fact, statistical indications of equality in gray matter volume in piriform cortex between groups could be demonstrated, a finding in sharp contrast to both the two previous studies in CA individuals as well as several studies assessing morphological effects in primary sensory areas due to life-long visual deprivation. Hence, the effect of lifelong absence of olfactory processing on this vital olfactory brain region is unclear.

Although scientists strive to continuously enhance the knowledge within their field of research, the scientific community struggle with what is often referred to as a reproducibility crisis, i.e., that published scientific results cannot be replicated and are therefore unlikely to reflect the ground truth (Baker, 2016; Open Science Collaboration, 2015). This can, at least partly, be accounted to the fact that the incentives for publishing new groundbreaking results often are stronger than the incentives for assessing whether new results reflect a true phenomenon (Smaldino and McElreath, 2016). The importance of further investigating unexpected results is particularly high when studying rare populations such as individuals with a congenital sensory deprivation, because these studies inherently deal with two major issues: limited sample sizes and between group comparisons. These issues generally lead to lower statistical power than wished for and increase the risk of, e.g., non-representative samples, factors that can lead to study outcomes that by chance do not reflect a phenomenon in the larger population sampled from; the aim of the majority of studies (Button et al., 2013).

We here aim to establish whether the piriform cortex is indeed morphologically unaffected by lifelong complete olfactory sensory deprivation and whether the morphological deviations are restricted to the OFC. To this end, we analyze the brain morphology of individuals with CA and normosmic controls, matched for age and sex, based on structural MRI. First, we investigate whether the unexpected results presented in Peter et al. (2020) are valid by doing a direct replication of the main analyses, i.e., assessing group differences with voxel- and surface-based morphometry. Second, to more thoroughly assess whether subtle patterns of group differences in gray matter volume exists, i.e., differences not detectable by the mass-univariate analyses applied in previous publications, we perform multivariate analysis using support vector machine (SVM) classification.

## 2. MATERIALS AND METHODS

To thoroughly assess whether the deviations in cerebral gray matter morphology in individuals with CA indeed is restricted to the OFC, the current study consists of two parts. First, a direct replication of our recent publication that demonstrated a surprising lack of morphological abnormalities in piriform cortex in individuals with CA compared to matched controls (Peter et al., 2020) was done using newly collected data (n_CA_=30, n_Control_=30;). Second, we combined the newly acquired dataset with a subset of the data from our previous publication (Peter et al., 2020)combined sample n_CA_=49, n_Control_=49, see details below) to examine whether subtle group differences in gray matter volume patterns exist using multivariate analysis in form of support vector machine (SVM) classification.

### 2.1 **Participants**

A total of 98 participants were included in this study: 49 individuals with isolated CA, i.e., congenital anosmia unrelated to specific genetic disorders, such as Kallmann syndrome, and 49 controls, matched in terms of sex and age (Table 1). Inclusion criteria for the CA group was a self-reported lifelong lack of olfactory perception without any known underlying condition causing the anosmia and a score in the anosmia range on the objective olfactory test (described below). For the control group, inclusion required a self-proclaimed and objectively verified functional sense of smell. Of the 49 CA individuals, 37 lacked bilateral olfactory bulbs, and the presence of bulbs in the remaining 12 was either non-determinable or identifiable but visibly hypoplastic (assessed from the structural images by J. N. L.).

**Table 1.**
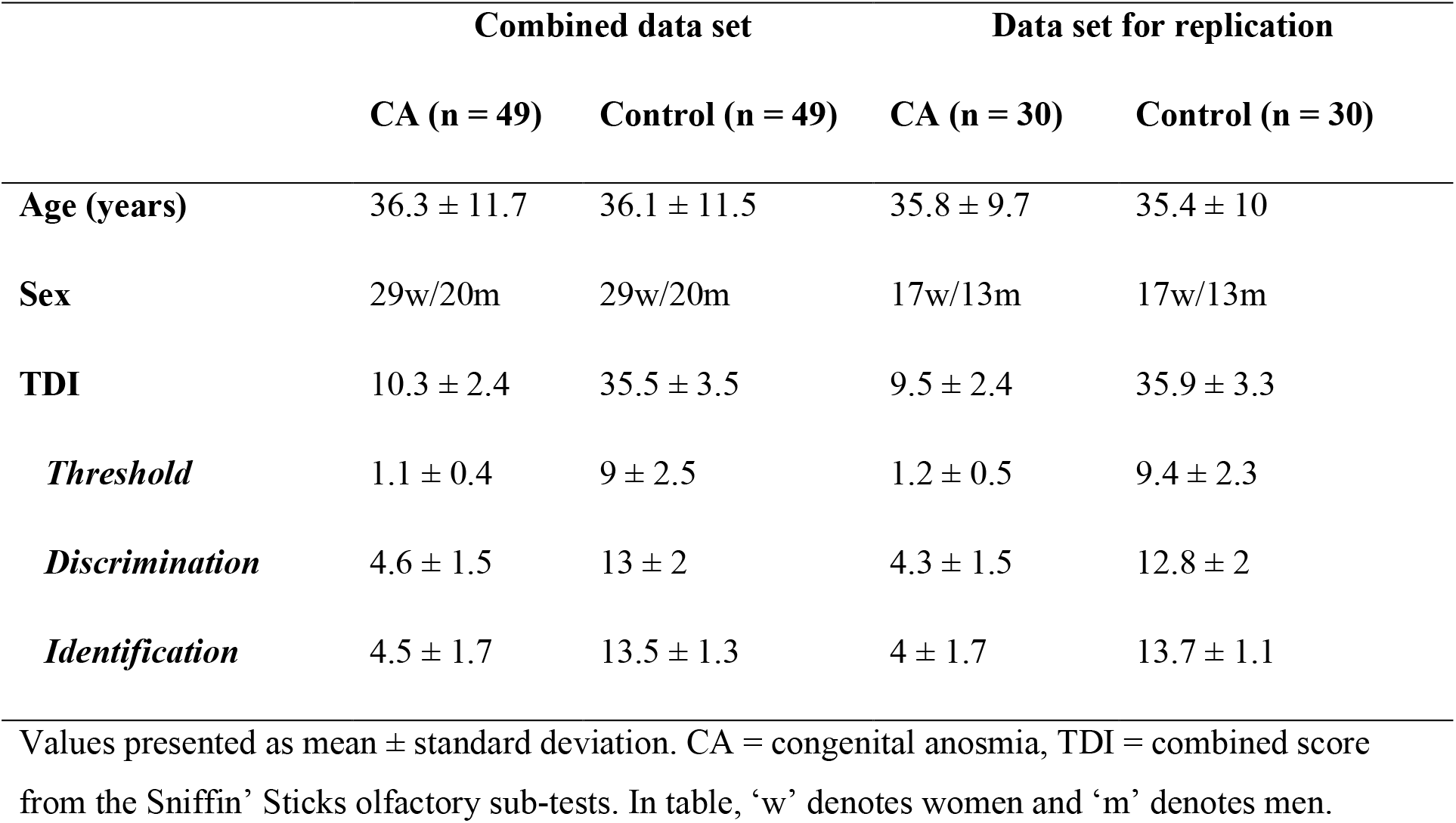
Descriptive statistics per experimental group.

|  | Combined data set |  | Data set for replication |  |
| --- | --- | --- | --- | --- |
|  | CA (n = 49) | Control (n = 49) | CA (n = 30) | Control (n = 30) |
| **Age (years)** | 36.3 ± 11.7 | 36.1 ± 11.5 | 35.8 ± 9.7 | 35.4 ± 10 |
| **Sex** | 29w/20m | 29w/20m | 17w/13m | 17w/13m |
| **TDI** | 10.3 ± 2.4 | 35.5 ± 3.5 | 9.5 ± 2.4 | 35.9 ± 3.3 |
| ***Threshold*** | 1.1 ± 0.4 | 9 ± 2.5 | 1.2 ± 0.5 | 9.4 ± 2.3 |
| ***Discrimination*** | 4.6 ± 1.5 | 13 ± 2 | 4.3 ± 1.5 | 12.8 ± 2 |
| ***Identification*** | 4.5 ± 1.7 | 13.5 ± 1.3 | 4 ± 1.7 | 13.7 ± 1.1 |

Values presented as mean ± standard deviation. CA = congenital anosmia, TDI = combined score from the Sniffin' Sticks olfactory sub-tests. In table, 'w' denotes women and 'm' denotes men.

The included data set consists of two parts: newly collected data and a subset of the data previously analyzed in Peter et al. (2020). The newly collected data set (30 individuals with CA, 30 matched controls) was acquired in Stockholm, Sweden, and used for the replication of Peter et al. (2020). Note that 13 of the individuals with CA in the newly collected data set had previously participated in Peter et al. (2020), but all controls were newly recruited. To optimize the conditions for the multivariate analysis by increasing the sample size, the newly collected data set was combined with data from all CA individuals from Peter et al. (2020) who did not participate in the new data collection, along with their matched controls (19 individuals with CA and 19 controls), resulting in the final set of 98 participants. This latter data was acquired at two sites: Wageningen, the Netherlands (11 individuals with CA, 11 matched controls) and Stockholm, Sweden (8 individuals with CA, 8 matched controls).

### 2.2 **Procedure**

#### 2.2.1 MR image acquisition

Siemens Magnetom 3T MR scanners (Siemens Healthcare, Erlangen, Germany) were used to acquire the imaging data: two Prisma scanners using 20-channel head coils (in Sweden) and a Verio scanner with a 32-channel head coil (in the Netherlands). T1-weighted images with whole-brain coverage were acquired using either a 3D GR/IR T1-weighted sequence (208 slices, TR = 2300 ms, TE = 2.89 ms, FA = 9°, voxel size = 1 mm^3^, FoV = 256 mm; 30 individuals with CA, 30 controls) or an MP-RAGE sequence (176 slices, TR = 1900 ms, TE = 2.52 ms, FA = 9°, voxel size = 1 mm^3^, FoV = 256 mm; 19 individuals with CA, 19 controls). Additional functional imaging data reported elsewhere were collected in the same testing sessions.

#### 2.2.2 Olfactory screening

To ensure functional anosmia and a normal sense of smell in respective groups, olfactory function of all participants was assessed with the full Sniffin’ Sticks olfactory test (Burghart, Wedel, Germany). The Sniffin’ Sticks test is a standardized test consisting of three subtests with individual scores: odor detection threshold (T), odor quality discrimination (D), and 4-alternative cued odor quality identification (I), together yielding the combined TDI-score used to classify individuals as normosmic, hyposmic, or anosmic based on normative data from over 9000 individuals (Oleszkiewicz et al., 2019). All individuals in the CA group were classified as functionally anosmic based on their TDI score and all individuals in the Control group were classified as normosmic within their age group (a score above the 10^th^ percentile in their group). Group mean scores on the Sniffin’ Stick olfactory performance test can be found in Table 1.

### 2.3 Replication

To assess whether the results presented in Peter et al. (2020) can be replicated, a direct replication of the main analyses was done using newly collected data: 30 individuals with CA and 30 matched controls (Table 1). Potential group differences in gray matter volume, cortical thickness, area, and curvature were estimated using mass-univariate analysis. To this end, the image processing and statistical analyses followed the analysis steps previously reported (Peter et al., 2020) using identical analysis scripts with the exception of the use of scanning site as a covariate in the analyses, because all participants were scanned at the same site in this newly acquired data set.

#### 2.3.1 Voxel-based morphometry

Voxel-based morphometry (VBM, Ashburner and Friston, 2000) analysis was done using SPM12 in MATLAB 2019b (The MathWorks, Inc., Natick, Massachusetts, USA). Segmentation of T_1_-weighted images into gray matter, white matter, and cerebrospinal fluid was done in native space using the unified segmentation approach (Ashburner and Friston, 2005). The three tissues were subsequently used to compute total intracranial volume for each participant and the gray and white matter were further used as input to a diffeomorphic image registration algorithm to improve inter-subject alignment (DARTEL, Ashburner, 2007). DARTEL aligns gray and white matter from all participants in an iterative process, creating an increasingly accurate average template for inter-subject alignment. The template and individual flow fields from DARTEL were used to spatially normalize gray matter images to MNI space with 12 parameter affine transformations. The normalized gray matter images were modulated with the Jacobian determinant of the deformation fields and smoothed with a 6 mm full-width at half maximum isotropic Gaussian kernel, with kernel size chosen to optimize the discovery of potential group differences in piriform cortex, a small region at the junction between frontal and temporal cortex.

To compensate for individual differences in intracranial volume when comparing gray matter volume between groups, the pre-processed, normalized gray matter images were proportionally scaled with total intracranial volume. Furthermore, the images were masked with a threshold of 0.15 to avoid inclusion of non-gray matter voxels. Voxel-wise group differences in gray matter volume were assessed using independent sample *t*-tests with age and sex as nuisance covariates. The tests were corrected for multiple comparisons with a family-wise error (FWE) corrected significance level of *p* < .05.

Following the analysis in Peter et al. (2020), in the absence of volumetric differences in piriform cortex from the voxel-wise approach, potential statistical evidence in support of a lack of group differences was investigated. To this end, individual mean gray matter volume from left and right piriform cortex were extracted using the MarsBaR toolbox (version 0.44, http://marsbar.sourceforge.net) based on a piriform cortex ROI from Zhou and colleagues (2019) and group differences were assessed based on Welch’s *t*-test. If the t-tests yielded non-significant results at an uncorrected significance level of *p* < .05, the two one-sided tests procedure for equivalence testing between groups was done using the TOSTER R package (Lakens, 2017) implemented in R (version 4.1.1, www.r-project.org). The equivalence bounds were defined based on the effect size that we can detect with 80% power (corresponding to *d* ≥ .65 based on 30 individuals in each group).

Because a subset of the CA individuals in this newly collected data set had previously participated in Peter et al. (2020), we wanted to ensure that potential replicated results were not driven exclusively by these individuals. Therefore, we conducted subgroup analyses of the CA individuals also included in Peter et al. (2020) (13 CA, 13 matched Controls) and the CA individuals not in Peter et al. (2020) (17 CA, 17 matched Controls) separately. Due to small sample sizes, a statistical threshold of *p* < .001 (uncorrected) was chosen to assess subgroup results.

#### 2.3.2 Surface based measures

The T_1_-weighted images were processed using FreeSurfer ver. 6.0 (Dale et al., 1999); http://surfer.nmr.mgh.harvard.edu/) to calculate cortical thickness, surface area, and underlying white matter curvature. FreeSurfer data was pre-processed through the HiveDB database system (Muehlboeck et al., 2014) and the image processing pipeline is described in detail in (Fischl and Dale, 2000). Processing steps include removal of non-brain tissue, Talairach transformation, segmentation of subcortical white matter and gray matter, intensity normalization, tessellation of the boundary between gray and white matter, automated topology correction, and surface deformation to find the boundary between gray and white matter as well as the boundary between gray matter and cerebrospinal fluid. Cortical thickness, surface area, and curvature were calculated at each vertex point on the tessellated surface in native space, after which individual surfaces were aligned to an average template and smoothed (5 mm full-width at half maximum surface based Gaussian kernel chosen to optimize analysis in piriform cortex) to enable statistical comparisons between the subject groups.

To assess group differences in cortical thickness, surface area, and curvature, vertex-wise values were compared between groups using a general linear model with age and sex as nuisance covariates. Correction for multiple comparisons was done based on a false discovery rate (FDR) of *p* < .05 and group differences surviving this threshold were defined as significant.

### 2.4 Support vector machine classification

After replication of the mass-univariate analyses of Peter et al. (2020), we examined whether potential fine-grained, wide-spread morphological differences between individuals with CA and normosmic controls exist, differences less likely to be detected by mass-univariate methods. To this end, we performed multivariate pattern analysis on a larger, combined data set consisting of the newly acquired data used for replication purposes and data from Peter et al. (2020). The CoSMoMVPA toolbox (https://www.cosmomvpa.org/) (Oosterhof et al., 2016) was used within MATLAB 2021b to investigated whether gray matter volume metrics derived from DARTEL (as described under the Voxel-based morphometry section) can be used as features to differentiate between individuals with CA and controls. To accomplish this, a searchlight approach was used to divide the smoothed gray matter images into spherical clusters with a radius of 6 mm. Subsequently, a linear support vector machine (SVM) was trained for each spherical cluster to classify whether each individual belongs to either the CA or control group. To select the training and testing groups, a cross-validation with 10-folds approach was performed to train the classifier on 80% of the subjects and test on the rest (randomly selected but balanced with respect to group). The group classification accuracy for each spherical cluster is projected on the center voxel of the sphere and the process is repeated for each sphere, resulting in a whole-brain classification accuracy map that illustrates how accurately subjects were classified into the CA and control groups. Voxels with classification accuracy > 75% were considered as significant. We then permuted the data 5000 times to test the statistical significance of the results. If the observed accuracy was more than 95% of the accuracies from permuted data, it was considered as significant (*p* < .05). (TFCE, Smith and Nichols, 2009)Thereafter, to specifically assess whether the gray matter volume patterns within piriform cortex differ between groups, i.e., could be used to discriminate between individuals with CA and controls, a directed analysis was performed using a ROI-based SVM classification. The gray matter volume values within bilateral piriform cortex were chosen as features and, in line with the searchlight analysis, cross-validation with 10-folds was performed with 80% for training and 20% for testing. Classification was repeated 5000 times on permuted data to assess the statistical significance of the results.

## 3. RESULTS

### 3.1 Replication of divergent morphological abnormalities within the orbitofrontal cortex in congenital anosmia

To explore CA-related alterations in gray matter volume, voxel-wise tests of group differences were performed. Four clusters demonstrated significant group differences, all located in the OFC: bilateral clusters of gray matter atrophy in the CA group along the olfactory sulci and bilateral clusters of gray matter volume increase in the CA group, centered around the medial orbital gyri (Figure 1B, Table 2). For visual comparison of the replication results, we also include the results of gray matter volumetric differences from Peter et. al., 2020 (Figure 1A) where corresponding clusters were found.

**Table 2.** Replication results from VBM-analysis.

| Region | Hemisphere | x<sup>a</sup> | y<sup>a</sup> | z<sup>a</sup> | Cluster size<sup>b</sup> | t<sup>c</sup> | p<sup>d</sup> |
| --- | --- | --- | --- | --- | --- | --- | --- |
| **CA > Control** |  |  |  |  |  |  |  |
| Medial orbital gyrus | L | -16 | 40 | -17 | 419 | 8.6 | < .001 |
| Medial orbital gyrus | R | 18 | 39 | -19 | 174 | 6.3 | .004 |
| **CA < Control** |  |  |  |  |  |  |  |
| Olfactory sulcus | L | -9 | 30 | -24 | 2686 | 11.1 | < .001 |
| Olfactory sulcus | R | 11 | 29 | -22 | 2898 | 11.0 | < .001 |

CA – congenital anosmia. <sup>a</sup> Coordinates derived from MNI152. <sup>b</sup> Cluster size (mm<sup>3</sup>). <sup>c</sup> t-value for peak voxel in cluster <sup>d</sup> p-value for peak voxel, FWE-corrected. Regions are identified based on ‘Atlas of the Human Brain’ (Mai et al., 2015).

**Figure 1.**
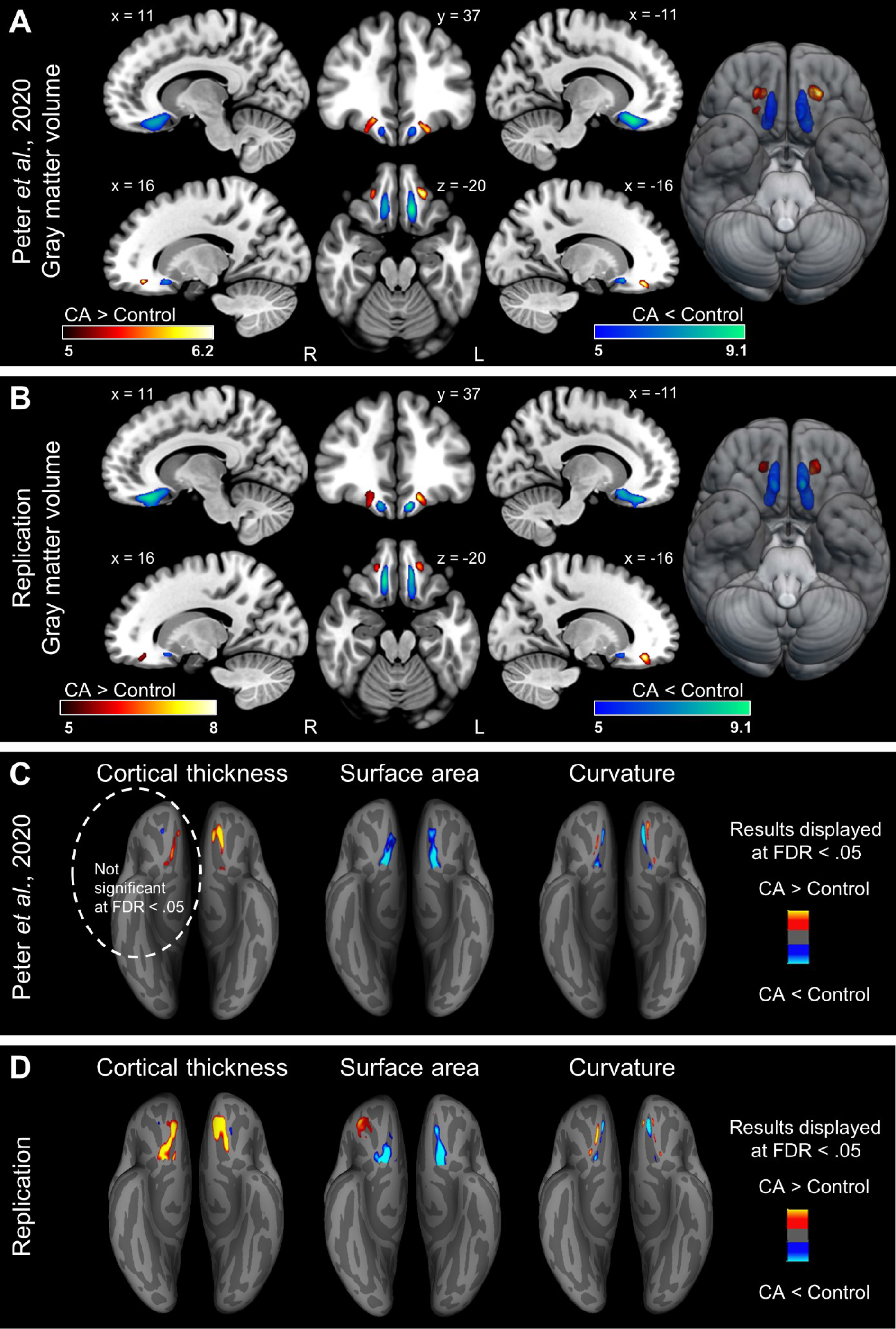
Results from replication. (**A**) Gray matter volumetric group differences recreated from the results previously reported in Peter et.al., 2020. (**B**) Gray matter volumetric group differences based on the current sample. Individuals with CA demonstrated gray matter atrophy in the bilateral olfactory sulci and gray matter volume increases in the bilateral medial orbital gyri. Results displayed at an FWE-corrected significance level of *p* < .05. (**C**) Group differences in surface-based measures previously presented in Peter et al., 2020. Note that group differences in cortical thickness in the right hemisphere are displayed at an uncorrected threshold of *p* < .001. (**D**) Group differences in surface-based measures based on the current sample. Results are displayed on an inferior view of an inflated brain at an FDR-corrected significance level of *p* < .05. Warm colors indicate larger values in the CA group compared to the Control group. Note specifically that for the curvature measure, all displayed results indicate decreases in absolute curvature in the CA group compared to the Control group because FreeSurfer’s convention of curvature is negative for gyri and positive for sulci. CA = congenital anosmia, R = right, L = left.

In line with our previous publication, no significant group differences in gray matter volume within piriform (primary olfactory) cortex were demonstrated at an FWE-corrected significance level of *p* < .05. To exclude the notion that the absence of differences in the piriform cortex is due to the conservative nature of the use of FWE-correction, we also assessed results using no correction for multiple comparisons and a threshold level of *p* < .001. Neither at this liberal threshold level did we observe any differences in piriform cortex (Supplementary Figure S1). Furthermore, no significant group differences in either right, *t*(57.7) = 0.39, *p* = .7, or left, *t*(58) = 1.05, *p* = .3, piriform cortex were demonstrated based on extraction of mean gray matter volume within our piriform ROI. We therefore performed equivalence testing using the two one-sided tests procedure to determine whether there is statistical evidence of an absence of group differences. There was statistically equivalent gray matter volume in both groups in the right piriform cortex (highest p-value: *t*(57.7) = 2.13, *p* = .019), i.e., supporting the conclusion that there is no difference in gray matter volume between the two groups in right piriform cortex. However, equivalence testing of the left piriform cortex did not lead to significant results based on the upper equivalence bound, *t*(58) = 1.47, *p* = .074. Consequently, the gray matter volume in the left piriform cortex could neither be deemed statistically different nor statistically equivalent between groups.

### 3.2 Major results consistent in both CA subgroups

The voxel-based morphometry replication results are near identical to the morphological group differences previously published in Peter et al., (2020, Figure 1A-B). To ensure that these similarities were not solely driven by the subgroup of CA individuals who participated in both data collections, two subgroup analyses were conducted: one with the CA individuals who had participated in the previous study and one with the CA individuals who had not. The major findings, i.e., the clusters of gray matter volume atrophy in the olfactory sulci and gray matter volume increases in the medial orbital gyri in individuals with CA compared to normosmic controls, were demonstrated in both CA subgroups (Supplementary Figure S2). It is furthermore noteworthy that no gray matter volume abnormalities in either left or right piriform cortex could be demonstrated in either CA subgroup based on Welch’s *t*-tests (all *p* > .377). Hence, we followed up with equivalence tests, resulting in statistical equivalence between the CA participants that were included in Peter *et al*. (2020) and matched controls in both left and right piriform cortex (largest *p* = .027). For the CA participants who did not participate in our previous study, statistical equivalence could be demonstrated in the right (largest *p* = .015) but not left (largest *p* = .055) piriform cortex. Full statistical results reported in Supplementary Table S1.

### 3.3 Thicker orbitofrontal cortex in the CA group

The volumetric analysis was followed up by surface-based analyses to assess potential group differences in cortical thickness, surface area, and curvature. Similar to the study here replicated (Peter et al., 2020), the demonstrated surface-based group differences were almost exclusively located in the OFC, as were the volumetric results. The cortical thickness analysis revealed a thickening of the bilateral medial orbital gyri and olfactory sulci in the CA group (Figure 1D, Table 3). The CA group further demonstrated decreased surface area in the bilateral olfactory sulci and increased surface area in the right, but not left, posterior orbital gyrus (Figure 1D, Table 3) accompanied by decreased curvature around the same orbitofrontal areas: the bilateral olfactory sulci and medial orbital gyri (Figure 1D, Table 3), all at an FDR threshold of < .05. Additional minor clusters with few vertices surviving our predefined threshold were identified and listed in Table 3, but not further addressed.

**Table 3.** Replication results from surface-based measures.

| Region | Hemisphere | x<sup>a</sup> | y<sup>a</sup> | z<sup>a</sup> | Cluster size<sup>b</sup> | p<sup>c</sup> |
| --- | --- | --- | --- | --- | --- | --- |
| **Cortical Thickness** |  |  |  |  |  |  |
| *CA > Control* |  |  |  |  |  |  |
| Olfactory sulcus, medial orbital gyrus | L | -8 | 38 | -22 | 338 | 10<sup>-15</sup> |
| Medial temporal gyrus | L | -47 | 12 | -26 | 12 | 5·10<sup>-5</sup> |
| Precentral gyrus | L | -36 | -29 | 52 | 15 | 7.9·10<sup>-5</sup> |
| Precentral gyrus, central sulcus | L | -35 | -22 | 49 | 4 | 1.6·10<sup>-4</sup> |
| Olfactory sulcus, medial orbital gyrus | R | 16 | 20 | -20 | 397 | 1.3·10<sup>-9</sup> |
| Olfactory sulcus, anterior olfactory nucleus | R | 10 | 12 | -16 | 27 | 2.5·10<sup>-6</sup> |
| Medial temporal gyrus | R | 51 | 11 | -20 | 11 | 1.3·10<sup>-5</sup> |
| Inferior frontal sulcus | R | 51 | 26 | 16 | 7 | 1.3·10<sup>-4</sup> |
| Medial temporal gyrus | R | 61 | -7 | -23 | 5 | 1.6·10<sup>-4</sup> |
| Fusiform gyrus | R | 33 | -5 | -38 | 1 | 2.5·10<sup>-4</sup> |
| *ICA < Control* |  |  |  |  |  |  |
| Medial orbital gyrus, posterior orbital sulcus | L | -19 | 31 | -17 | 18 | 2.5·10<sup>-5</sup> |
| Medial orbital gyrus | R | 23 | 35 | -15 | 9 | 1.3·10<sup>-4</sup> |
| **Area** |  |  |  |  |  |  |
| *CA > Control* |  |  |  |  |  |  |
| Gyrus rectus | L | -6 | 47 | -22 | 2 | 1.3·10<sup>-4</sup> |
| Posterior orbital gyrus, transverse orbital sulcus | R | 37 | 37 | -8 | 280 | 2.5·10<sup>-6</sup> |
| Posterior orbital gyrus | R | 32 | 28 | -18 | 4 | 3.2·10<sup>-4</sup> |
| *CA < Control* |  |  |  |  |  |  |
| Olfactory sulcus | L | -13 | 11 | -15 | 251 | 2.5·10<sup>-10</sup> |
| Olfactory sulcus | R | 14 | 16 | -13 | 256 | 1.6·10<sup>-10</sup> |
| Olfactory sulcus | R | 13 | 33 | -18 | 19 | 2·10<sup>-4</sup> |
| Gyrus rectus | R | 7 | 13 | -14 | 4 | 1.6·10<sup>-4</sup> |
| **Curvature** |  |  |  |  |  |  |
| *CA < Control<sup>d</sup>* |  |  |  |  |  |  |
| Olfactory sulcus | L | -10 | 40 | -20 | 59 | 7.9·10<sup>-12</sup> |
| Medial orbital gyrus | L | -14 | 39 | -22 | 13 | 2.5·10<sup>-6</sup> |
| Posterior orbital gyrus | L | -17 | 15 | -22 | 11 | 3.2·10<sup>-6</sup> |
| Gyrus rectus | L | -4 | 41 | -24 | 9 | $7.9 \cdot 10^{-8}$ |
| Olfactory sulcus | L | -16 | 12 | -15 | 7 | $4 \cdot 10^{-5}$ |
| Posterior angular gyrus | L | -35 | -75 | 42 | 6 | $1.6 \cdot 10^{-5}$ |
| Medial orbital gyrus | L | -15 | 29 | -22 | 6 | $2 \cdot 10^{-5}$ |
| Medial orbital gyrus | R | 14 | 27 | -25 | 40 | $10^{-8}$ |
| Olfactory sulcus | R | 13 | 39 | -20 | 32 | $4 \cdot 10^{-8}$ |
| Posterior orbital sulcus | R | 18 | 12 | -15 | 29 | $6.3 \cdot 10^{-7}$ |
| Posterior orbital gyrus | R | 19 | 14 | -22 | 14 | $7.9 \cdot 10^{-7}$ |
| Olfactory sulcus | R | 15 | 29 | -19 | 9 | $7.9 \cdot 10^{-6}$ |

### 3.4 SVM correctly classified CA and Controls in the olfactory sulci but not piriform cortex

Both voxel-wise and vertex-wise (i.e., surface-based) analyses revealed consistent morphological differences between CA and control individuals in the OFC. We subsequently aimed to determine whether a multivariate method in form of SVM classification could differentiate between these two groups based on gray matter volumes, specifically in the piriform cortex. The multivariate analysis, thresholded at an accuracy level of 75%, revealed one significant cluster, centered around the bilateral olfactory sulci, that distinguished between CA individuals and normosmic controls based on their gray matter volume (cluster size 28641 mm^3^, peak coordinate: x −14, y 35, z −27, peak accuracy: 95.3%, *p* = .009, Figure 2). Although the vast majority of piriform cortex was not included in the cluster that allowed the groups to be differentiated, there was a slight overlap between the posterior end of the significant cluster and the anterior piriform cortex (Supplementary Figure S3). We therefore performed more specific ROI-based analysis solely on the piriform cortex. The gray matter volume metrics in piriform cortex alone were not sufficient to accurately classify participants as belonging to the CA and control groups (classification accuracy 52.5%, *p* = .33).

**Figure 2.**
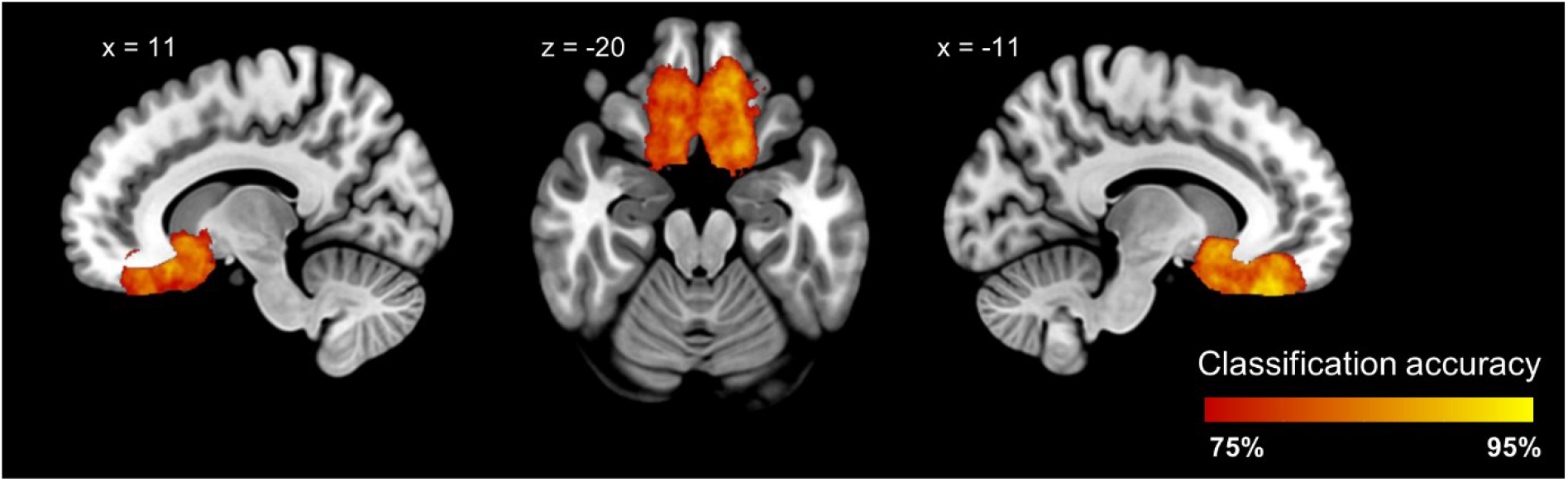
Group classification accuracy based on gray matter volume. Significant cluster located around the bilateral olfactory sulci with classification accuracy > 75% (*p* = .009).

## 4. DISCUSSION

It has been clearly established that lifelong visual sensory deprivation alters the morphology of visual cortical areas. Whether a lifelong absence of olfactory experiences alters the morphology of the human olfactory cortex has been under debate with studies yielding diverging results. We here demonstrate that individuals with isolated congenital anosmia (CA) have seemingly normal morphology in piriform (olfactory) cortex, thus confirming the surprising results presented in Peter et al. (2020). Further, we demonstrate bilateral clusters of morphological alterations in the orbitofrontal cortex (OFC) in individuals with CA compared to matched healthy controls, result that line up well with the existing literature (Frasnelli et al., 2013; Karstensen et al., 2018; Peter et al., 2020). These results are obtained both in a direct replication of the mass-univariate analysis from Peter et al. (2020) and with multivariate analysis methods.

The seemingly normal morphology in piriform cortex in individuals with CA demonstrated in Peter et al. (2020) contradicted the results from two previous studies on CA morphology (Frasnelli et al., 2013; Karstensen et al., 2018) as well as studies on individuals with acquired anosmia (Bitter et al., 2010; Iravani et al., 2021a; Peng et al., 2013; Yao et al., 2014; but see Yao et al., 2018). In the current study, we set out to establish whether the results obtained in Peter et al. (2020) accurately reflect the brain morphology in CA. In a direct replication of Peter et al. (2020) we can here demonstrate near identical results to those previously presented, namely an absence of group differences in piriform cortex as well as bilateral clusters of morphological group differences in the OFC: decreases around the olfactory sulci and increases around the medial orbital gyri for the CA group. We further demonstrate that the replication results are not driven by an overlap of participants in the two studies. First, a new group of matched controls was included in the current study. This ensures that the demonstrated group differences are not driven by a randomly selected unrepresentative control group. This phenomenon is often overlooked, but was clearly demonstrated in a study where the tactile performance of blind individuals was enhanced only in comparison to one of the two included sighted control groups (Alary et al., 2009). Second, we show in a subgroup analysis that all major results were reproduced in both the sample of CA individuals who participated in both the recent data collection and in Peter et al. (2020) (n = 13) as well as those unique to the current study (n = 17). Note that when assessing the subgroups, a more liberal statistical threshold was used due to the smaller sample sizes, resulting in additional small clusters of group differences that lack overlap between the two subgroups; these results could have been interpreted as interesting findings in a context where information from larger samples was missing. This puts greater emphasis on the importance of increasing statistical power and replicating studies to establish trustworthiness of findings in research fields where small sample sizes is common.

In addition to replicating the major results from the original study (Peter et al., 2020), we can here demonstrate bilateral clusters of increased cortical thickness around the olfactory sulci, whereas the cluster in the right hemisphere failed to reach significance when correcting for multiple comparisons in Peter et al. (2020). The bilateral result presented here is likely related to lower variability as a result of all data being collected at one site, using one scanner and head coil, whereas two scanning sites were used to increase the sample size in Peter et al. (2020). Still, the lower variability did not lead to detection of any group differences in piriform cortex, neither in a whole-brain analysis correcting for multiple comparisons, nor in a directed ROI-analysis. In fact, equivalence of piriform gray matter volume between groups was demonstrated unilaterally, whereas the gray matter volume was not statistically different but not statistically equal unilaterally (Lakens et al., 2018). Hence, by replicating the analyses in (Peter et al., 2020), we can here in a new data set confirm the morphological deviations in the OFC and, importantly, the lack thereof in piriform cortex in individuals with CA.

Mass-univariate analyses, as those discussed above, are the most frequently used analyses in the study of brain morphology in sensory deprivation and have contributed with much knowledge. Mass-univariate analyses are, however, by nature limited to detect large, localized effects. In contrast, multivariate analysis methods enable the detection of more complex and subtle patterns (Davatzikos, 2019) and therefore have the potential to reveal gray matter volume abnormalities in individuals with CA neither detected in Peter et al. (2020) nor in the current replication. Hence, to complement the mass-univariate analyses applied in previous studies of CA (Frasnelli et al., 2013; Karstensen et al., 2018; Peter et al., 2020), the recently collected data was combined with data from Peter et al. (2020) and analyzed with support vector machine (SVM) classification. Using a searchlight approach, which enables localized classification accuracy results, one large cluster with high (>75%) classification accuracy located in the bilateral OFC was identified, i.e., covering the regions also identified with the mass-univariate analyses. This overlap of results between methods is expected – although the SVM has the potential to detect more widespread, subtle patterns of group differences, it will also utilize the larger, more localized effects that the mass-univariate methods can detect. The significant cluster from the SVM classification connected the previously identified regions and extended beyond them, with the posterior part reaching towards the anterior piriform cortex and into the caudate nucleus. An enlargement of clusters is a feature of the searchlight analysis approach that needs to be taken into account when interpreting the results. It stems from the fact that the classification accuracy projected on one voxel is based on all information within a sphere surrounding the voxel, meaning that the same voxel contributing to high classification accuracy will be included in all spheres at a distance less than or equal to the sphere radius (here 6mm) from that voxel. Hence, because the few voxels in piriform cortex showing high classification accuracy are located at the posterior end of the large orbitofrontal cluster, it is likely that they are merely a projection of the group differences in the orbitofrontal cortex. This interpretation is supported by the fact that an SVM classifier was unable to distinguish between the two groups when only the voxels within piriform cortex were used as features for classification in a targeted analysis. Although a multivariate analysis approach arguably is better suited than mass-univariate analyses when assessing brain morphology in olfactory sensory deprivation based on the heterogeneity of the olfactory system (Iravani et al., 2021b), the classification results in the current study line up well with the mass-univariate results and, importantly, does not identify additional brain regions of group differences. Hence, also the multivariate analyses provide strong support for large clusters of gray matter volumetric group differences and that these group differences are restricted to the OFC.

Individuals with CA demonstrate a lack of indications of morphological abnormalities in piriform cortex, a cortical region of outmost importance for olfactory processing (Gottfried, 2006; Lundström et al., 2011; Seubert et al., 2013), independent of whether uni- or multivariate analysis methods are applied. The question that remains to be answered is why the morphology of this region is seemingly unaffected by a lifelong absence of the expected input whereas visual cortex demonstrates clear morphological reorganization when deprived of visual input (Jiang et al., 2015; Lee and Whitt, 2015; Noppeney, 2007; Voss, 2019). Atrophy in piriform cortex within individuals with acquired anosmia has been demonstrated (Bitter et al., 2010; Iravani et al., 2021a; Peng et al., 2013; Yao et al., 2014; but see Yao et al., 2018), thereby indicating that the structure of the region is plastic and can be affected by altered input. A potential explanation to our lack of effects might be that the morphology in piriform cortex is preserved in individuals with CA precisely because the olfactory sensory deprivation is congenital and not later acquired, thereby not experiencing a sudden change in sensory input and a subsequent push to restore homeostasis via plastic reorganization (Pascual-Leone et al., 2005). That a clear shift in sensory input might trigger plasticity is supported by findings in animal models where the cortical thickness of piriform cortex is seemingly unaffected by a deprivation of olfactory input right after birth. In contrast, olfactory bulb ablation later in life leads to noticeable decrease in piriform cortex thickness (Friedman and Price, 1986; Westrum and Bakay, 1986). A competing hypothesis is that the absence of (macroscopic) morphological deviations in piriform cortex demonstrated here could be related to a functional overtake where the piriform cortex processes non-olfactory information. Indeed, piriform activity has been shown to be elicited by non-olfactory input (Gottfried et al., 2004; Porada et al., 2019) in individuals with a normal sense of smell, suggesting that cross-modal processing might be present also in individuals with CA. These studies do, however, explicitly or implicitly associate the presented non-olfactory stimuli with olfactory stimuli and it is therefore unknown whether demonstrated cross-modal activity is dependent on olfactory association or not, an association that by nature is non-existing in individuals with CA.

In clear contrast to piriform cortex, considerable morphological deviations are demonstrated in olfactory related areas of the OFC in individuals with CA. Of particular interest is the fact that these morphological deviations seem to be specific to congenital anosmia: when compared to acquired olfactory deprivation of different etiologies, the demonstrated results bore a striking resemblance to the results our current and previous study, where individuals with CA are compared to normosmic controls (Postma et al., 2021). Following the speculations of Peter et al. (2020), the volumetric decreases around the olfactory sulci are likely related to the absence or hypoplasia of the olfactory bulbs distinguishing individuals with CA given that the olfactory bulbs are located right below the olfactory sulci. Thus, it can be assumed that these group differences are based on a congenital morphological abnormality in individuals with CA rather than a plastic rearrangement of cortex as an effect of absent olfactory input (Peter et al., 2020). In contrast, the morphological increases demonstrated in the OFC overlap quite well with the orbitofrontal subregion demonstrating the highest likelihood for olfactory-induced activity based on neuroimaging studies (Seubert et al., 2013). This implies that these morphological abnormalities are a likely consequence of the absence of olfactory processing in that region, either based on decreased synaptic pruning due to absent olfactory input (Frasnelli et al., 2013), related to compensatory processing from our other senses (Merabet and Pascual-Leone, 2010), or both. Indeed, we recently showed that individuals with CA demonstrated enhanced audio-visual integration performance compared to healthy controls (Peter et al., 2019). Note, however, that no group differences in audio-visual processing in the OFC were found in a subsequent imaging study comparing individuals with CA and matched controls (Peter et al., 2021).

A strength of the current study is the large sample size, in the context of studying congenital sensory deprivation and other rare populations. The size of our data set do, however, still limit the statistical power, i.e., decreasing the probability of finding small effects if present in the sample as well as increasing the probability that detected effects in fact are spurious (Button et al., 2013). The fact that the morphological groups differences detected in OFC and the absence of differences in piriform cortex are supported independent of applied analysis method supports both the findings and null findings. It should, however, be noted that the highly interesting null-finding in piriform cortex does not necessarily imply that the region is completely structurally normal in individuals with a lifelong absence of olfactory processing. We can here only provide evidence for a lack of gross abnormalities given the type of data and analysis methods used; we are thus also limited by the spatial resolution of the structural images (1 mm^3^) and cannot dismiss the possibility that group differences in the neuronal constitution and cortical layering exists.

We can here conclusively demonstrate that lifelong olfactory deprivation creates large morphological deviations in gray matter within areas of the OFC where CA individuals experience both areas of increase and areas of decrease. Critically, we can replicate and more firmly demonstrate that lifelong olfactory sensory deprivation does not alter the gross anatomical organization of the piriform cortex, a region argued to constitute primary olfactory cortex. Why the primary olfactory cortex, in sharp contrast to other sensory cortices, is not affected by long-term sensory deprivation and what functional and behavioral effects this results in remains to be determined. Taken together, our results suggest that the olfactory system is not affected by sensory deprivation in the same manner as our other sensory systems: lifelong olfactory sensory deprivation mainly affects higher-order olfactory processing regions.

## CONFLICT OF INTEREST

The authors report no conflict of interest.

## ACKNOWLEDGEMENTS

This work was supported by the Knut and Alice Wallenberg Foundation (KAW 2018.0152) and the Swedish Research Council (2017-02325; 2021-02700), all awarded to J.N.L.. The use of the MR facility at Stockholm University (SUBIC) was made possible by a grant to SUBIC (SU FV-5.1.2-1035-15). The use of the 3T MRI facility at Hospital Gelderse Vallei in Ede (NL) has been made possible by Wageningen University & Research Shared Research Facilities.

---

**Table S1.**
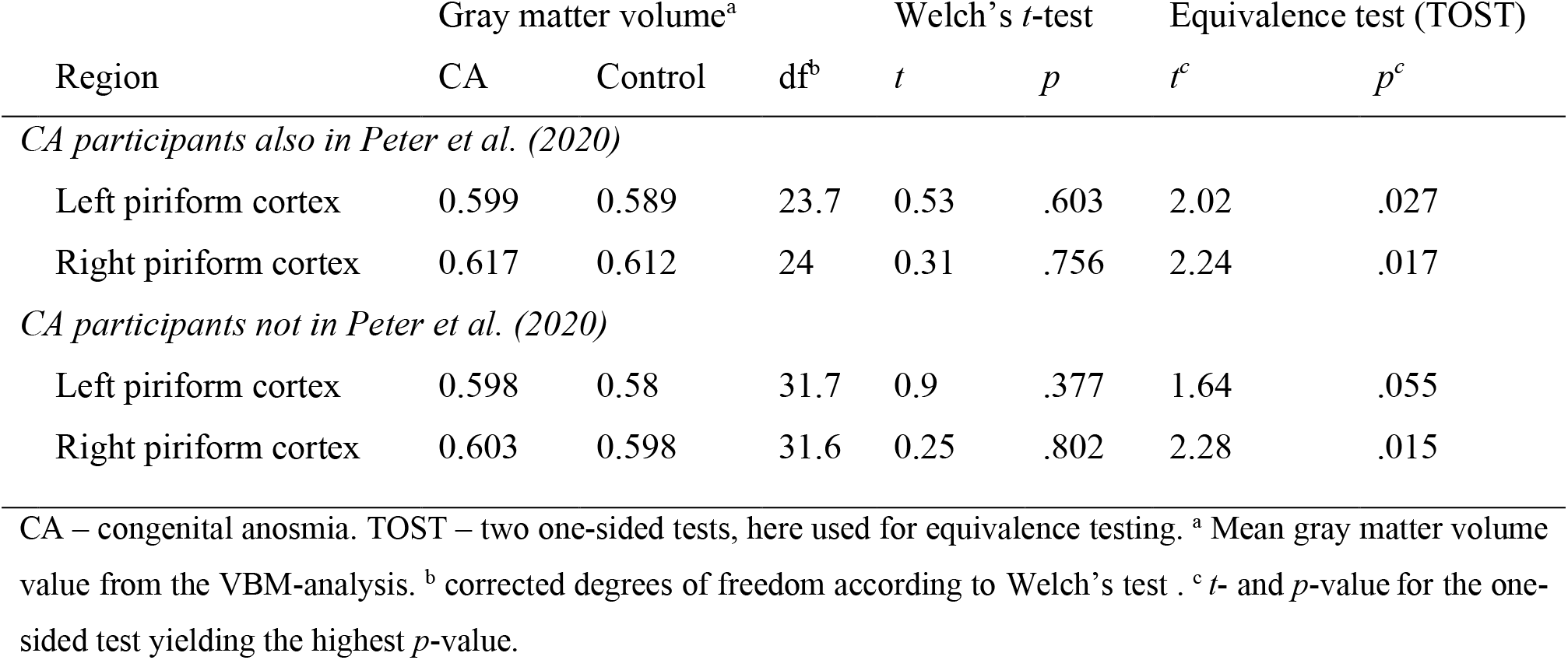
Subgroup analysis piriform cortex

**Supplementary Figure S1.**
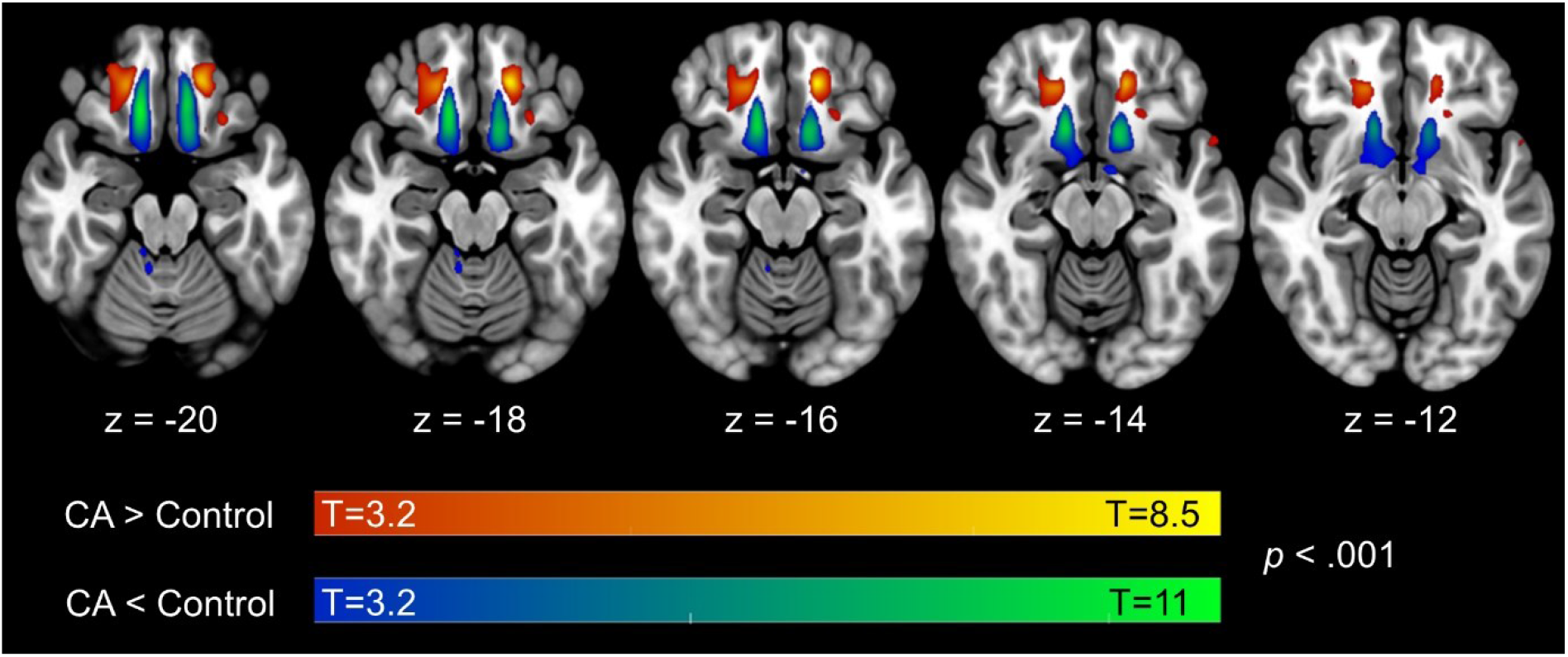
Replication results of group differences in gray matter volume. displayed on axial slices thresholded at an uncorrected significance level of p < .001. No indications of volumetric group differences in piriform cortex demonstrated at this uncorrected significance level.

**Supplementary Figure S2.**
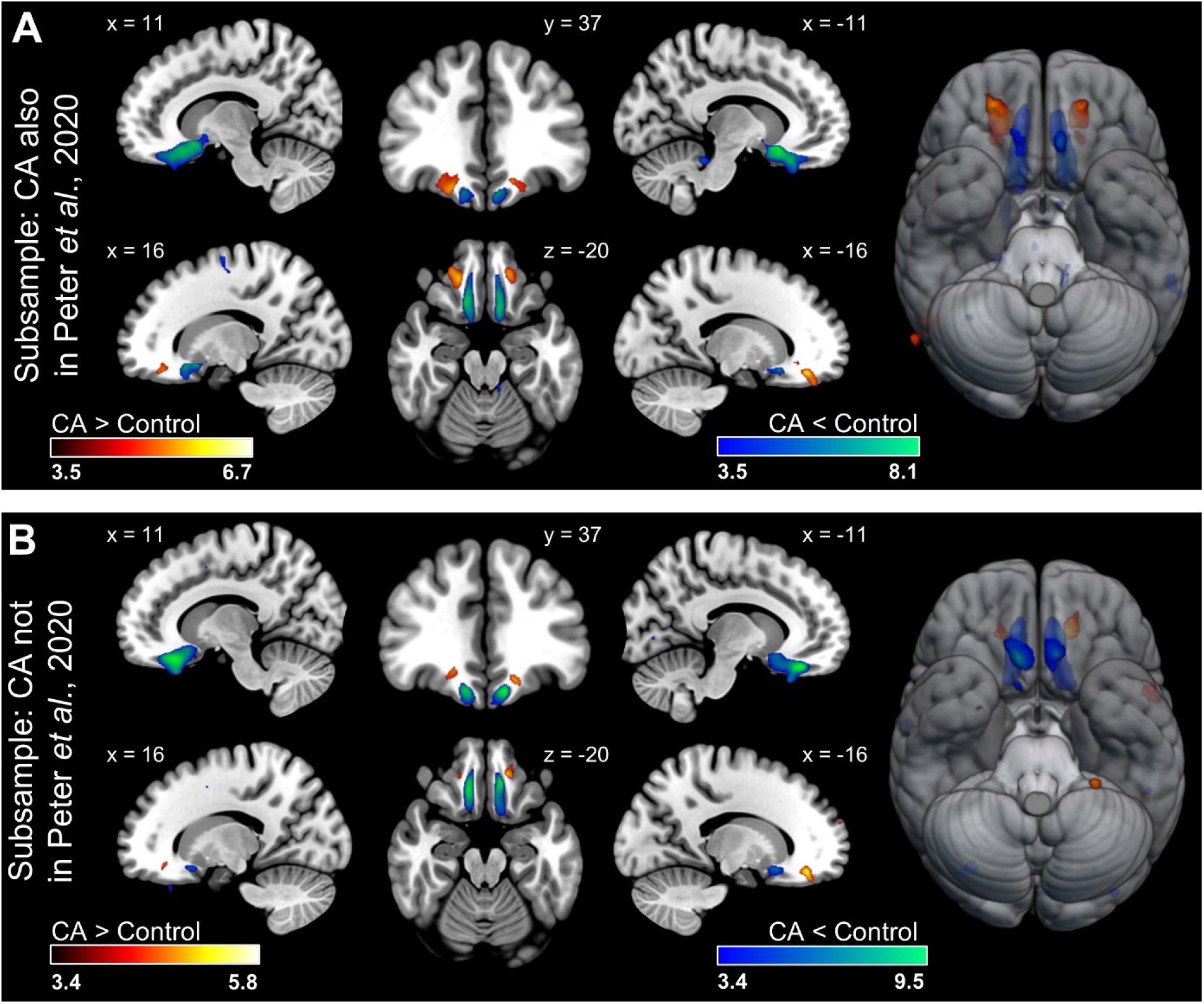
Replication results of group differences in gray matter volume for the two CA subgroups and matched controls. **A)** CA participants who also participated in Peter et al. (2020) compared to matched controls (N_CA_ = 13, N_Control_ = 13). **B)** CA participants who did not participate in Peter et al. (2020) compared to matched controls (N_CA_ = 17, N_Control_ = 17). Both subgroups of individuals with CA demonstrated gray matter atrophy in the bilateral olfactory sulci and gray matter volume increases in the bilateral medial orbital gyri. Results displayed at an uncorrected significance level of *p* < .001.

**Supplementary Figure S3.**
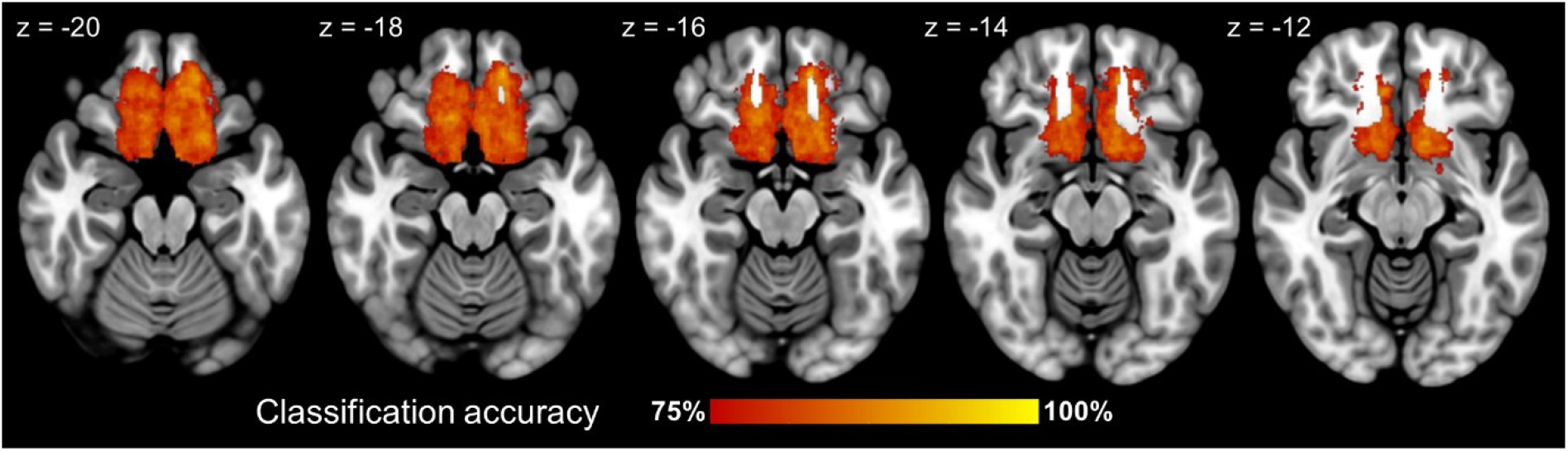
Group classification accuracy based on gray matter volume.

## Notes

### Competing Interest Statement

The authors have declared no competing interest.

## REFERENCES

Alary, F., Duquette, M., Goldstein, R., Elaine Chapman, C., Voss, P., La Buissonnière-Ariza, V., Lepore, F., 2009. Tactile acuity in the blind: a closer look reveals superiority over the sighted in some but not all cutaneous tasks. Neuropsychologia 47, 2037–2043. doi:10.1016/j.neuropsychologia.2009.03.014

Ashburner, J., 2007. A fast diffeomorphic image registration algorithm. Neuroimage 38, 95–113. doi:10.1016/j.neuroimage.2007.07.007

Ashburner, J., Friston, K.J., 2000. Voxel-based morphometry--the methods. Neuroimage 11, 805–821. doi:10.1006/nimg.2000.0582

Ashburner, J., Friston, K.J., 2005. Unified segmentation. Neuroimage 26, 839–851. doi:10.1016/j.neuroimage.2005.02.018

Baker, M., 2016. 1,500 scientists lift the lid on reproducibility. Nature 533, 452–454. doi:10.1038/533452a

Bitter, T., Gudziol, H., Burmeister, H.P., Mentzel, H.-J., Guntinas-Lichius, O., Gaser, C., 2010. Anosmia leads to a loss of gray matter in cortical brain areas. Chem. Senses 35, 407–415. doi:10.1093/chemse/bjq028

Boesveldt, S., Postma, E.M., Boak, D., Welge-Luessen, A., Schöpf, V., Mainland, J.D., Martens, J., Ngai, J., Duffy, V.B., 2017. Anosmia-A Clinical Review. Chem. Senses 42, 513–523. doi:10.1093/chemse/bjx025

Button, K.S., Ioannidis, J.P.A., Mokrysz, C., Nosek, B.A., Flint, J., Robinson, E.S.J., Munafò, M.R., 2013. Power failure: why small sample size undermines the reliability of neuroscience. Nat. Rev. Neurosci. 14, 365–376. doi:10.1038/nrn3475

Dale, A.M., Fischl, B., Sereno, M.I., 1999. Cortical surface-based analysis. I. Segmentation and surface reconstruction. Neuroimage 9, 179–194. doi:10.1006/nimg.1998.0395

Davatzikos, C., 2019. Machine learning in neuroimaging: Progress and challenges. Neuroimage 197, 652–656. doi:10.1016/j.neuroimage.2018.10.003

Fischl, B., Dale, A.M., 2000. Measuring the thickness of the human cerebral cortex from magnetic resonance images. Proc. Natl. Acad. Sci. USA 97, 11050–11055. doi:10.1073/pnas.200033797

Frasnelli, J., Fark, T., Lehmann, J., Gerber, J., Hummel, T., 2013. Brain structure is changed in congenital anosmia. Neuroimage 83, 1074–1080. doi:10.1016/j.neuroimage.2013.07.070

Freiherr, J., 2017. Cortical Olfactory Processing, in: Buettner, A. (Ed.), Springer handbook of odor. Springer International Publishing, Cham, pp. 97–98. doi:10.1007/978-3-319-26932-0_38

Friedman, B., Price, J.L., 1986. Plasticity in the olfactory cortex: age-dependent effects of deafferentation. J. Comp. Neurol. 246, 1–19. doi:10.1002/cne.902460102

Gottfried, J.A., 2006. Smell: central nervous processing. Adv. Otorhinolaryngol. 63, 44–69. doi:10.1159/000093750

Gottfried, J.A., 2010. Central mechanisms of odour object perception. Nat. Rev. Neurosci. 11, 628–641. doi:10.1038/nrn2883

Gottfried, J.A., Smith, A.P.R., Rugg, M.D., Dolan, R.J., 2004. Remembrance of odors past: human olfactory cortex in cross-modal recognition memory. Neuron 42, 687–695. doi:10.1016/s0896-6273(04)00270-3

Gottfried, J.A., Zald, D.H., 2005. On the scent of human olfactory orbitofrontal cortex: meta-analysis and comparison to non-human primates. Brain Res Brain Res Rev 50, 287–304. doi:10.1016/j.brainresrev.2005.08.004

Haberly, L.B., 2001. Parallel-distributed processing in olfactory cortex: new insights from morphological and physiological analysis of neuronal circuitry. Chem. Senses 26, 551–576. doi:10.1093/chemse/26.5.551

Iravani, B., Peter, M.G., Arshamian, A., Olsson, M.J., Hummel, T., Kitzler, H.H., Lundström, J.N., 2021a. Acquired olfactory loss alters functional connectivity and morphology. BioRxiv. doi:10.1101/2021.01.11.426175

Iravani, B., Peter, M.G., Arshamian, A., Olsson, M.J., Hummel, T., Kitzler, H.H., Lundström, J.N., 2021b. Acquired olfactory loss alters functional connectivity and morphology. Sci. Rep. 11, 16422. doi:10.1038/s41598-021-95968-7

Jiang, A., Tian, J., Li, R., Liu, Y., Jiang, T., Qin, W., Yu, C., 2015. Alterations of regional spontaneous brain activity and gray matter volume in the blind. Neural Plast. 2015, 141950. doi:10.1155/2015/141950

Jiang, J., Zhu, W., Shi, F., Liu, Y., Li, J., Qin, W., Li, K., Yu, C., Jiang, T., 2009. Thick visual cortex in the early blind. J. Neurosci. 29, 2205–2211. doi:10.1523/JNEUROSCI.5451-08.2009

Karstensen, H.G., Vestergaard, M., Baaré, W.F.C., Skimminge, A., Djurhuus, B., Ellefsen, B., Brüggemann, N., Klausen, C., Leffers, A.-M., Tommerup, N., Siebner, H.R., 2018. Congenital olfactory impairment is linked to cortical changes in prefrontal and limbic brain regions. Brain Imaging Behav. 12, 1569–1582. doi:10.1007/s11682-017-9817-5

Lakens, D., 2017. Equivalence Tests: A Practical Primer for t Tests, Correlations, and Meta-Analyses. Soc. Psychol. Personal. Sci. 8, 355–362. doi:10.1177/1948550617697177

Lakens, D., Scheel, A.M., Isager, P.M., 2018. Equivalence testing for psychological research: A tutorial. Advances in Methods and Practices in Psychological Science 1, 259–269. doi:10.1177/2515245918770963

Lee, H.-K., Whitt, J.L., 2015. Cross-modal synaptic plasticity in adult primary sensory cortices. Curr. Opin. Neurobiol. 35, 119–126. doi:10.1016/j.conb.2015.08.002

Lundström, J.N., Boesveldt, S., Albrecht, J., 2011. Central processing of the chemical senses: an overview. ACS Chem. Neurosci. 2, 5–16. doi:10.1021/cn1000843

Mai, J., Majtanik, M., Paxinos, G., 2015. Atlas of the Human Brain, 4th Edition. ed.

Manan, H.A., Yahya, N., Han, P., Hummel, T., 2022. A systematic review of olfactory-related brain structural changes in patients with congenital or acquired anosmia. Brain Struct. Funct. 227, 177–202. doi:10.1007/s00429-021-02397-3

Merabet, L.B., Pascual-Leone, A., 2010. Neural reorganization following sensory loss: the opportunity of change. Nat. Rev. Neurosci. 11, 44–52. doi:10.1038/nrn2758

Muehlboeck, J.-S., Westman, E., Simmons, A., 2014. TheHiveDB image data management and analysis framework. Front Neuroinformatics 7, 49. doi:10.3389/fninf.2013.00049

Noppeney, U., 2007. The effects of visual deprivation on functional and structural organization of the human brain. Neurosci. Biobehav. Rev. 31, 1169–1180. doi:10.1016/j.neubiorev.2007.04.012

Noppeney, U., Friston, K.J., Ashburner, J., Frackowiak, R., Price, C.J., 2005. Early visual deprivation induces structural plasticity in gray and white matter. Curr. Biol. 15, R488–R490. doi:10.1016/j.cub.2005.06.053

Oleszkiewicz, A., Schriever, V.A., Croy, I., Hähner, A., Hummel, T., 2019. Updated Sniffin’ Sticks normative data based on an extended sample of 9139 subjects. Eur. Arch. Otorhinolaryngol. 276, 719–728. doi:10.1007/s00405-018-5248-1

Oosterhof, N.N., Connolly, A.C., Haxby, J.V., 2016. CoSMoMVPA: Multi-Modal Multivariate Pattern Analysis of Neuroimaging Data in Matlab/GNU Octave. Front Neuroinformatics 10, 27. doi:10.3389/fninf.2016.00027

Open Science Collaboration, 2015. Estimating the reproducibility of psychological science. Science 349, aac4716. doi:10.1126/science.aac4716

Park, H.-J., Lee, J.D., Kim, E.Y., Park, B., Oh, M.-K., Lee, S., Kim, J.-J., 2009. Morphological alterations in the congenital blind based on the analysis of cortical thickness and surface area. Neuroimage 47, 98–106. doi:10.1016/j.neuroimage.2009.03.076

Pascual-Leone, A., Amedi, A., Fregni, F., Merabet, L.B., 2005. The plastic human brain cortex. Annu. Rev. Neurosci. 28, 377–401. doi:10.1146/annurev.neuro.27.070203.144216

Peng, P., Gu, H., Xiao, W., Si, L.F., Wang, J.F., Wang, S.K., Zhai, R.Y., Wei, Y.X., 2013. A voxel-based morphometry study of anosmic patients. Br. J. Radiol. 86, 20130207. doi:10.1259/bjr.20130207

Peter, M.G., Mårtensson, G., Postma, E.M., Engström Nordin, L., Westman, E., Boesveldt, S., Lundström, J.N., 2021. Seeing Beyond Your Nose? The Effects of Lifelong Olfactory Sensory Deprivation on Cerebral Audio-visual Integration. Neuroscience 472, 1–10. doi:10.1016/j.neuroscience.2021.07.017

Peter, M.G., Mårtensson, G., Postma, E.M., Nordin, L.E., Westman, E., Boesveldt, S., Lundström, J.N., 2020. Morphological changes in secondary, but not primary, sensory cortex in individuals with life-long olfactory sensory deprivation. Neuroimage 218, 117005. doi:10.1016/j.neuroimage.2020.117005

Peter, M.G., Porada, D.K., Regenbogen, C., Olsson, M.J., Lundström, J.N., 2019. Sensory loss enhances multisensory integration performance. Cortex 120, 116–130. doi:10.1016/j.cortex.2019.06.003

Porada, D.K., Regenbogen, C., Seubert, J., Freiherr, J., Lundström, J.N., 2019. Multisensory enhancement of odor object processing in primary olfactory cortex. Neuroscience 418, 254–265. doi:10.1016/j.neuroscience.2019.08.040

Postma, E.M., Smeets, P.A.M., Boek, W.M., Boesveldt, S., 2021. Investigating morphological changes in the brain in relation to etiology and duration of olfactory dysfunction with voxel-based morphometry. Sci. Rep. 11, 12704. doi:10.1038/s41598-021-92224-w

Ptito, M., Schneider, F.C.G., Paulson, O.B., Kupers, R., 2008. Alterations of the visual pathways in congenital blindness. Exp. Brain Res. 187, 41–49. doi:10.1007/s00221-008-1273-4

Reichert, J.L., Schöpf, V., 2018. Olfactory loss and regain: lessons for neuroplasticity. Neuroscientist 24, 22–35. doi:10.1177/1073858417703910

Seubert, J., Freiherr, J., Djordjevic, J., Lundström, J.N., 2013. Statistical localization of human olfactory cortex. Neuroimage 66, 333–342. doi:10.1016/j.neuroimage.2012.10.030

Smaldino, P.E., McElreath, R., 2016. The natural selection of bad science. R. Soc. Open Sci. 3, 160384. doi:10.1098/rsos.160384

Smith, S.M., Nichols, T.E., 2009. Threshold-free cluster enhancement: addressing problems of smoothing, threshold dependence and localisation in cluster inference. Neuroimage 44, 83–98. doi:10.1016/j.neuroimage.2008.03.061

Voss, P., 2019. Brain (re)organization following visual loss. Wiley Interdiscip Rev Cogn Sci 10, e1468. doi:10.1002/wcs.1468

Westrum, L.E., Bakay, R.A., 1986. Plasticity in the rat olfactory cortex. J. Comp. Neurol. 243, 195–206. doi:10.1002/cne.902430205

Yao, L., Pinto, J.M., Yi, X., Li, L., Peng, P., Wei, Y., 2014. Gray matter volume reduction of olfactory cortices in patients with idiopathic olfactory loss. Chem. Senses 39, 755–760. doi:10.1093/chemse/bju047

Yao, L., Yi, X., Pinto, J.M., Yuan, X., Guo, Y., Liu, Y., Wei, Y., 2018. Olfactory cortex and Olfactory bulb volume alterations in patients with post-infectious Olfactory loss. Brain Imaging Behav. 12, 1355–1362. doi:10.1007/s11682-017-9807-7

Zelano, C., Bensafi, M., Porter, J., Mainland, J., Johnson, B., Bremner, E., Telles, C., Khan, R., Sobel, N., 2005. Attentional modulation in human primary olfactory cortex. Nat. Neurosci. 8, 114–120. doi:10.1038/nn1368

Zelano, C., Mohanty, A., Gottfried, J.A., 2011. Olfactory predictive codes and stimulus templates in piriform cortex. Neuron 72, 178–187. doi:10.1016/j.neuron.2011.08.010

Zhou, G., Lane, G., Cooper, S.L., Kahnt, T., Zelano, C., 2019. Characterizing functional pathways of the human olfactory system. Elife 8. doi:10.7554/eLife.47177

